# Dynamically controlled light delivery over large brain volumes through tapered optical fibers

**DOI:** 10.1101/094524

**Authors:** Ferruccio Pisanello, Gil Mandelbaum, Marco Pisanello, Ian A. Oldenburg, Leonardo Sileo, Jeffrey E. Markowitz, Ralph E. Peterson, Andrea Della Patria, Trevor M. Haynes, Mohamed S. Emara, Barbara Spagnolo, Sandeep R. Datta, Bernardo L. Sabatini, Massimo De Vittorio

## Abstract

Optogenetics promises spatiotemporal precise control of neural processes using light. However, the spatial extent of illumination within the brain is difficult to control and cannot be adjusted using standard fiber optics. We demonstrate that optical fibers with tapered tips can be used to illuminate either large brain volumes or dynamically selectable subregions. Remotely adjusting the light input angle to the fiber varies the light-emitting portion of the taper over several millimeters without movement of the implant. We use this mode to activate dorsal versus ventral striatum of individual mice and reveal different effects of each manipulation on motor behavior. Conversely injecting light over the full numerical aperture of the fiber results in light emission from the entire taper surface, achieving broader and more efficient optogenetic activation of neurons when compared to the standard flat-faced fiber stimulation. Thus, tapered fibers permit focal or broad illumination that can be precisely and dynamically matched to experimental needs.

## INTRODUCTION

Optogenetic modulation of neuronal activity has become the dominant method of examining the behavioral consequences of activity in specific neuronal populations *in vivo*. The prevalence of experimental paradigms based on optogenetics is due to the synergy of research advances in two distinct but well-connected fields: the development of ever-improving light-activated modulators of electrical activity ^1–3^ and of technologies to deliver light within the brain of free-moving animals ^4–8^. Nevertheless, attaining the full potential of optical neural control requires new technologies to better control the spatial extent of light delivery and more precisely match illumination to the heterogeneous structure of the brain.

In certain applications, it is necessary to deliver uniform illumination to large brain areas, whereas for other applications confined illumination of small brain volumes is preferred. Ideally, both modes of illumination could be accomplished via a single, reconfigurable device. To this aim a number of approaches have been developed, including multiple implanted waveguides^8–10^, multiple micro light delivery devices (μLEDs)^6,7,11,12^ and multi-point emitting optical fibers (MPFs)^13,14^. All these techniques allow for site selective light delivery, and can extend the illuminated brain volume by activating multiple emission points. Furthermore, the light delivery pattern can be reconfigured during the experiment. On the other hand, they are limited by several downsides: implanting multiple waveguides is highly invasive, μLEDs can heat tissue during prolonged illumination, and MPFs require higher input laser power to produce viable optogenetic control. Importantly, likely due to the difficulty and cost of building the required devices, these approaches remain niche applications and have not been broadly utilized in neuroscience labs. Indeed, the most common light delivery method for optogenetic experiments remains flat-faced optical fibers (FFs), which deliver highly spatially heterogeneous illumination to a relatively small and fixed brain volume near the fiber facet. Furthermore, due to the relatively large and flat area of the cleaved end, these fibers can cause substantial tissue damage during insertion.

Here we describe a tapered optical fiber (TF) whose emission properties can be simply and dynamically reconfigured to switch between relatively homogenous large volume light delivery and spatially restricted illumination. Multiple wavelengths of light can be independently modulated and directed to subvolumes of interest. The device consists of a single, thin and sharp waveguide, thus minimizing its invasiveness. To demonstrate the suitability of this approach for more uniform and efficient illumination of extended brain structures, experiments were performed in the primary motor cortex and striatum of awake head-restrained and freely moving mice. We also demonstrate that by controlling the angle at which light is injected into the fiber, TFs can emit light from sub-portions of the taper to produce spatio-temporally resolved patterns that subsample the volume of interest. We use this approach to achieve site specific optogenetic stimulation and demonstrate that activation of indirect pathway striatal projection neurons (iSPNs) in dorsal vs. ventral striatum has different effects on locomotion in freely moving mice exploring an open arena. Thus TFs provide a simple, inexpensive, and easy to operate multipurpose system for optical control of neural activity.

## RESULTS

### Design principles of tapered optical fibers

TFs are multimode fiber optics that have been engineered to taper gradually from their full width (125-225 μm) to ~500 nm. The taper angle is small (2°<ψ<8°) such that the taper length varies between 1.5-5.6 mm (Fig. 1). This design was chosen to permit smooth insertion into the brain, reduce the implant cross-section, and expose a large area of the fiber core for potential light emission. Three-dimensional ray-tracing and geometric models demonstrate the working principle of the device (Fig. 1a-b). A ray injected into the core/cladding section of the fiber with an input angle θ is guided via total internal reflection (TIR) to the tapered region. At each reflection of the ray its propagation angle with respect to the fiber optical axis increases by an amount equal to the taper angle ψ (Fig. S1). This occurs until a critical section is met, at which TIR is lost and the ray therefore radiates into the surrounding medium. The distance between this light-emission point and the taper tip depends on θ such that increasing θ moves the emission section further from the tip (Fig. 1b). The dependency of the output from the taper tip on input angle is determined by the fiber numerical aperture (NA) and taper angle (Fig. S2 for optical fibers with NA 0.22 and 0.39 and ψ=2.2° and ψ=2.9°, respectively). Therefore, the light input angle θ selects the output zone along the length of a specific taper.

**Figure 1.**
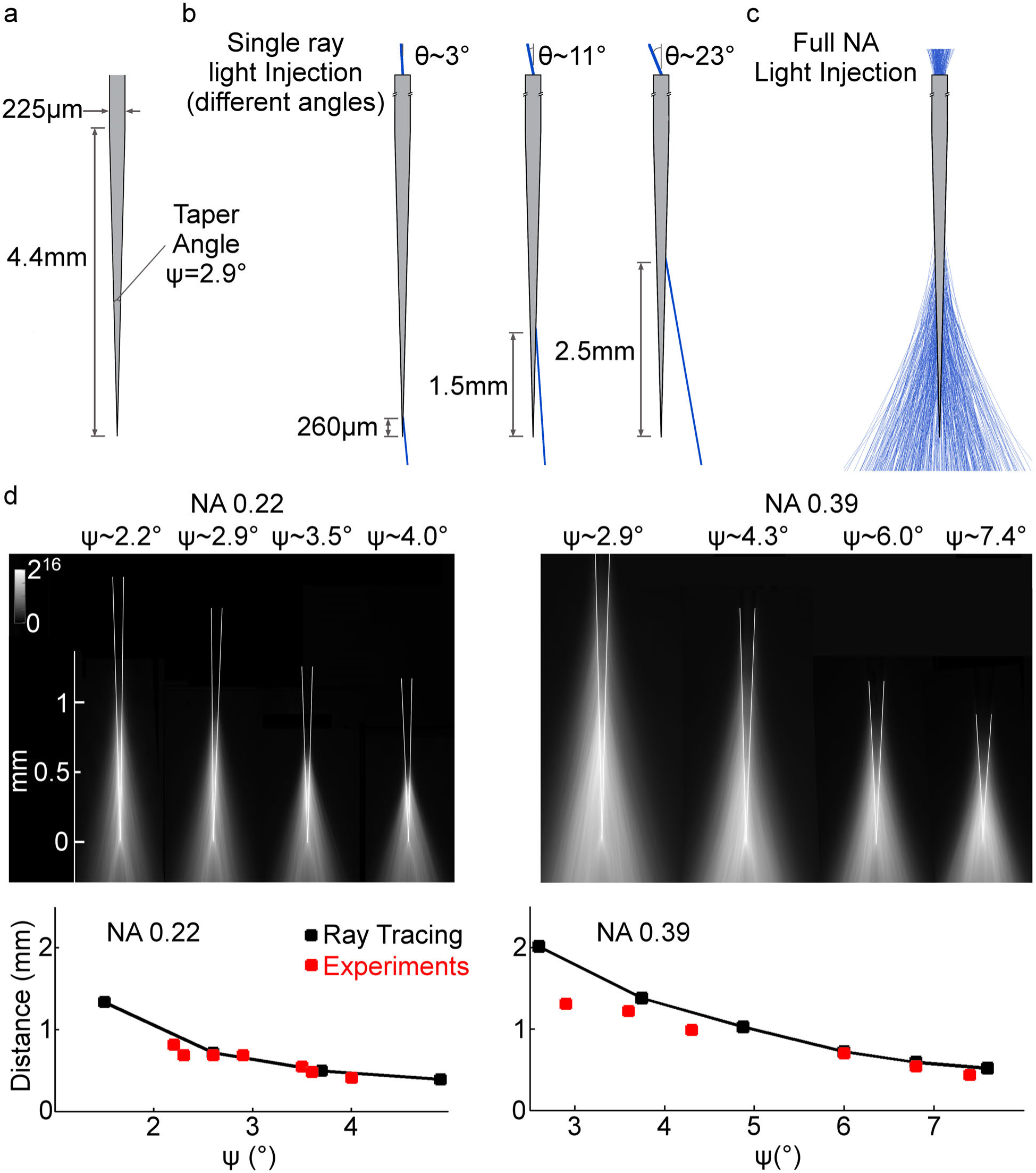
Emission properties of TFs. **a**, Schematic representation of a typical TF geometry (NA=0.39, taper angle ψ=2.9°, taper length 4.4 mm, core/cladding diameters 200/225 μm). **b**, Ray tracing simulations of emission from the taper tip resulting from injecting a single ray in the fiber at different angles. **c**, Ray distributions resulting from injecting light using the full NA of the fiber. **d**, *top*, Image of fluorescence generated by light emitted from tapered fibers with the specified geometries immersed in a fluorescein solution. *bottom*, The graphs depict calculated (black) and measured (red) emission lengths (evaluated as the full width at half maximum *L_0.5_*) for 0.22 NA and 0.39 NA TFs as a function of the taper angle. 0.22 NA TFs have core/cladding diameters of 50/125 μm, whereas 0.39 NA TFs have core/cladding diameters of 200/225 μm.

A direct consequence of this dependence of the position of light emission on θ is that TFs can be used to emit light along the fiber in two fundamentally different ways. First, when the full NA at the fiber input is used to inject light into the TF, light is emitted from a broad extent of the taper (Fig. 1c), as desired for illumination of spatially extended brain regions. For example, the entire cortical thickness or the dorsal-ventral access of the striatum. For a fiber of a particular NA, the length of the light-emitting segment (*L*) depends mainly on the taper angle. Ray tracing simulations indicate that *L* can be tailored from a few hundred micrometers up to a few millimeters in the case of 0.22 and 0.39 NA fibers with taper angles ψ ranging from 2.2° to 7.4° (Fig. S3).

We experimentally verified this effect in 0.22 and 0.39 NA fibers of various taper angles by immersing the taper in a fluorescein solution and imaging the resulting fluorescence distribution (Fig. S4). A decrease of the taper angle increases the length of the emitting segment up to ~1 mm in the case of 0.22 NA 50 μm/125 μm core/cladding fibers and up to ~2 mm in the case of 0.39 NA 200 μm/225 μm fibers (Fig. 1d). Thus, in this light-injection mode, a TF with the proper NA and taper angle can be chosen to match the linear extent of light output to the size of many mouse brain structures.

In order to compare the ray-tracing model with experimental results, we evaluated the emission length, referred to as *L_0.5_*, over which the delivered intensity is maintained above 50% of its peak (Fig. S4). We found good agreement between modeling and experiments for both 0.39 NA and 0.22 NA fibers (Fig. 1d and Fig. S4). The slight differences observed at low ψ for 0.39 NA fibers arise because the taper is assumed to be perfectly linear in the ray tracing model, whereas the real structure has a modest parabolic shape (Fig. S5). The diameter at which light starts to out-couple and the total light power delivered are nearly independent of the taper angle (Figs. S6). As a consequence, manufacturing TFs with lower taper angle ψ spreads the available power over a larger taper surface (Fig. S7 and Fig. S8), potentially allowing light delivery to a more elongated brain region.

### Illumination of large brain volumes with TFs

Flat cleaved optical fibers (‘Flat Fibers’ or FFs) are commonly inserted just above a brain volume of interest and the delivered light is strongly attenuated by the tissue, allowing excitation of neurons located only up to a few hundreds of micrometers from the fiber end^15^ ^17^. Significant excitation of neurons further from the fiber face requires large increases in laser power to overcome the typical exponential decay in power density from the fiber face. In contrast, by virtue of their thin and sharp edge, TFs can be inserted into the volume of interest and light delivered along the length of the taper (Fig. 2). To evaluate the illumination pattern achieved in light absorbing and scattering brain tissue, we implanted TFs and FFs into fluorescein impregnated fixed mouse brain slices and imaged the fluorescence generated by light emitted from the fiber (Fig. 2a). FFs illuminate a small brain volume and fluorescence is strongly attenuated after a few hundred micrometers from the emitting flat end facet (Fig. 2b). In contrast, TFs emit light along the taper length, resulting in elongated and more homogenous illumination of the tissue (Fig. 2c–d).

**Figure 2.**
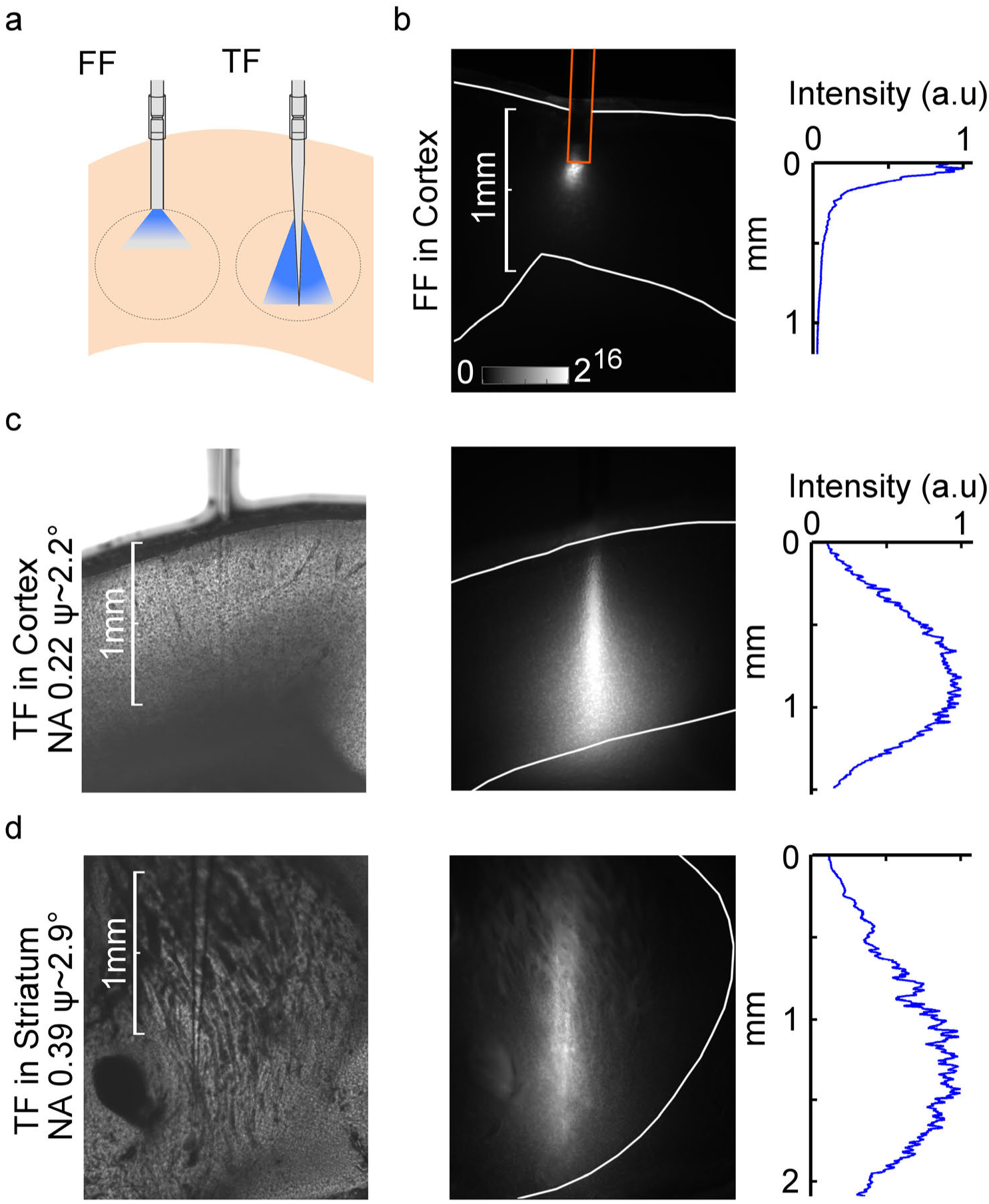
Emission properties of TFs in brain slices. **a**, Schematic of light delivery in brain tissue through FFs and TFs. **b**, *left*, Image of fluorescence induced by light emission from an FF implanted into cortex in a fluorescein impregnated brain slice. Gray scale represents fluorescence intensity in arbitrary units of a linear scale. *right*, Normalized fluorescence intensity profile in the tissue starting from the fiber end face **c** and **d**, Bright field images (*left*) to identify the position of the TFs in the fluorescein impregnated cortical or striatal brain slice used for acquisition of the fluorescence images (*middle*). Gray scale is the same as in (b). *right*, Normalized profiles of fluorescence intensities beside the taper, starting from the first emission point.

The differences in tissue illumination achieved by TF and FF arise from two important differences: (1) TFs emit light along a larger surface, i.e. the cone defined by L and ψ; (2) Light emerges from the TFs at a non-zero angle with respect to the taper axis (e.g. Fig. 1c). As a consequence, tissue absorption and scattering do not determine light distribution along the fiber axis as in FFs, but along the direction of emitted light, which has a significant component perpendicular to the taper axis. Notably, the depth of the excited volume can be tailored by selecting the taper geometry and the fiber NA, instead of by increasing the laser power as commonly done in experiments with FFs. For instance, TFs with 0.22 NA and ψ=2.2° deliver light for the whole cortical depth, whereas using TFs with 0.39 NA and ψ=2.9° most of the depth of the striatum can be illuminated (Fig. 2c–d).

### *In vivo* examination of effective excitation in striatum, a large brain structure

In order to characterize the activity patterns associated with TFs light delivery *in vivo*, we compared the ability of TFs and FFs to activate ChR2 expressing cells in striatum, a large subcortical nucleus. Either a TF (0.39 NA and ψ=2.9°) or a FF (0.22 NA or 0.39 NA) was implanted in the striatum of transgenic mice *(Ador2a-Cre; Ai32)* expressing ChR2 in the indirect striatal projection (iSPNs) neurons (Fig. 3). This mouse was selected for analysis because iSPNs are GABAergic neurons that locally inhibit other striatal neurons and inhibit recurrent excitatory inputs into striatum, minimizing secondary activation of cells not expressing ChR2 ^18^. The TF was implanted at a depth of 3.7 mm and the FF at 2.3 mm, respectively. Light (473 nm, 1 mW outputted at fiber exit) was delivered at a 30 sec on/30 sec off cycle for 1 hour to awake animals in their home cages. To compare the spatial distribution of cells activated by light delivered through TFs and FFs, animals were euthanized 2 hours post stimulation and we performed fluorescence immunohistochemistry for c-fos, the protein product of an immediate early gene whose expression is regulated by neuronal activity ^19^. Stimulation through the TF resulted in more uniform c-fos induction across approximately 2 mm of the dorsal ventral axis of the striatum (Fig. 3a–b), compared to the more spatially restricted induction in FF implanted animals (Fig. 3c–d). Furthermore, although light delivery through the TF stimulated cells throughout the dorsal/ventral axis (i.e. along the axis of the fiber), differential placement of the fiber permitted selective stimulation of either lateral or medial sub-regions of striatum (Fig. 3b). Thus, as suggested by the simulations and fluorescence excitation *ex vivo* (Figs. 1–2), *in vivo* TFs deliver light across a spatially extended volume of tissue surrounding the thin fiber. Control experiments showed minimal c-fos in animals that expressed ChR2 in iSPNs but did not receive light stimulation and in wild type animals negative for ChR2 that did receive light stimulation (Fig. S9). Lastly, a clear advantage of the TF compared to FF was the minimal damage induced in the tissue (Fig. S10), likely a result of the tapered profile that culminates in a sub-micron size tip.

**Figure 3.**
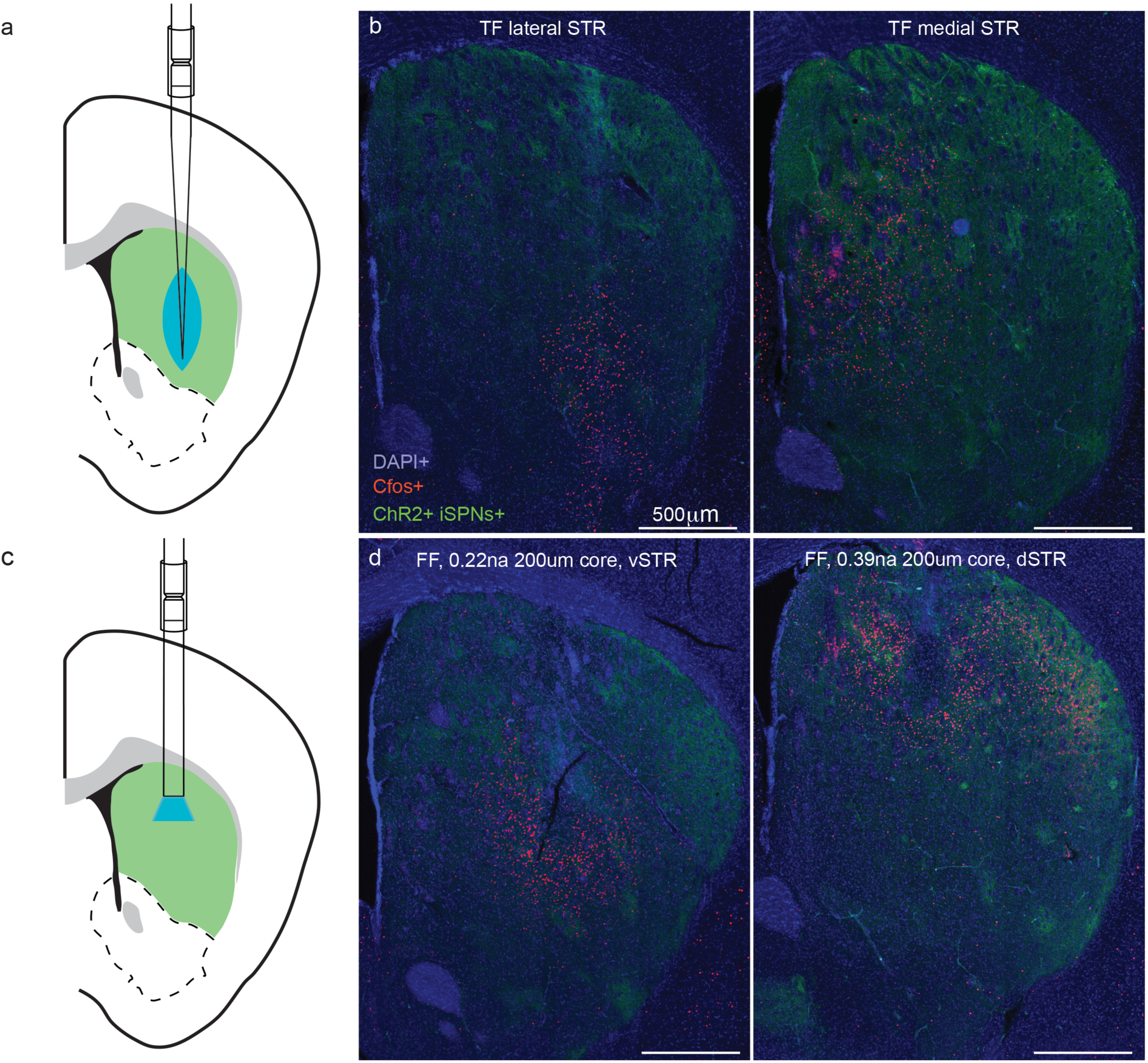
*In vivo* examination of effective excitation in striatum. **a**, Schematic of experimental preparation showing a TF inserted into the striatum of a mouse expressing ChR2 in indirect pathway SPNs (iSPNs). **b**, c-fos expression (red) in the striatum (coronal section, 0.85 mm anterior to Bregma) of an animal expressing ChR2-YFP in iSPNs (green) after light stimulation delivered by a TF in lateral (*left*) or medial (*right*) striatum. DAPI is shown in blue. **c**, Schematic of experimental preparation as in (a) showing placement of a flat-faced fiber (FF). **d**, As in (b) showing c-fos induction by light delivery from a FF in dorsal (*left*, 0.22 NA) or ventral (*right*, 0.39 NA) striatum.

### Optogenetic control of motor cortex with TFs

In order to examine the potential benefits of more uniform light delivery *in vivo*, TFs were tested in the primary motor cortex of awake head-restrained *VGAT-ChR2 BAC* transgenic mouse ^20^, which express ChR2 in all inhibitory neurons (Fig. 4a). A TF (NA 0.22, ψ*~*2.2°) and a FF were implanted serially near a 16-contact silicon multi-electrode array (Neuronexus). The FF was placed such that the flat cleaved face was at the same depth of the first emission point (shallow position) or the tip (deep position) of the TF. Light (473 nm) was delivered as 5 x 50 ms pulses at 5 Hz, repeated every 3 s for 80 times. Output powers from the TF and FF were matched in order to examine the efficiency of each to inhibit cortex via stimulation of GABAergic interneurons. TFs more effective suppressed cortical activity at lower power levels (Fig. 4b and Fig. S11): Inhibition with TFs was obtained at ~10 μW of total power (~0.20 mW/mm^2^, see Fig. S8 for estimated power density distribution along the taper) whereas FF (NA=0.22, core diameter 50 μm) required ~5-fold higher power to obtain a comparable effect at both depths (50 μW total output power, corresponding to ~25 mW/mm^2^, according to ref. [^21^]). Moreover, even at higher powers, inhibition is more pronounced with TFs, suggesting that tapered fibers stimulate a higher number of ChR2-expressing GABAergic neurons.

**Figure 4.**
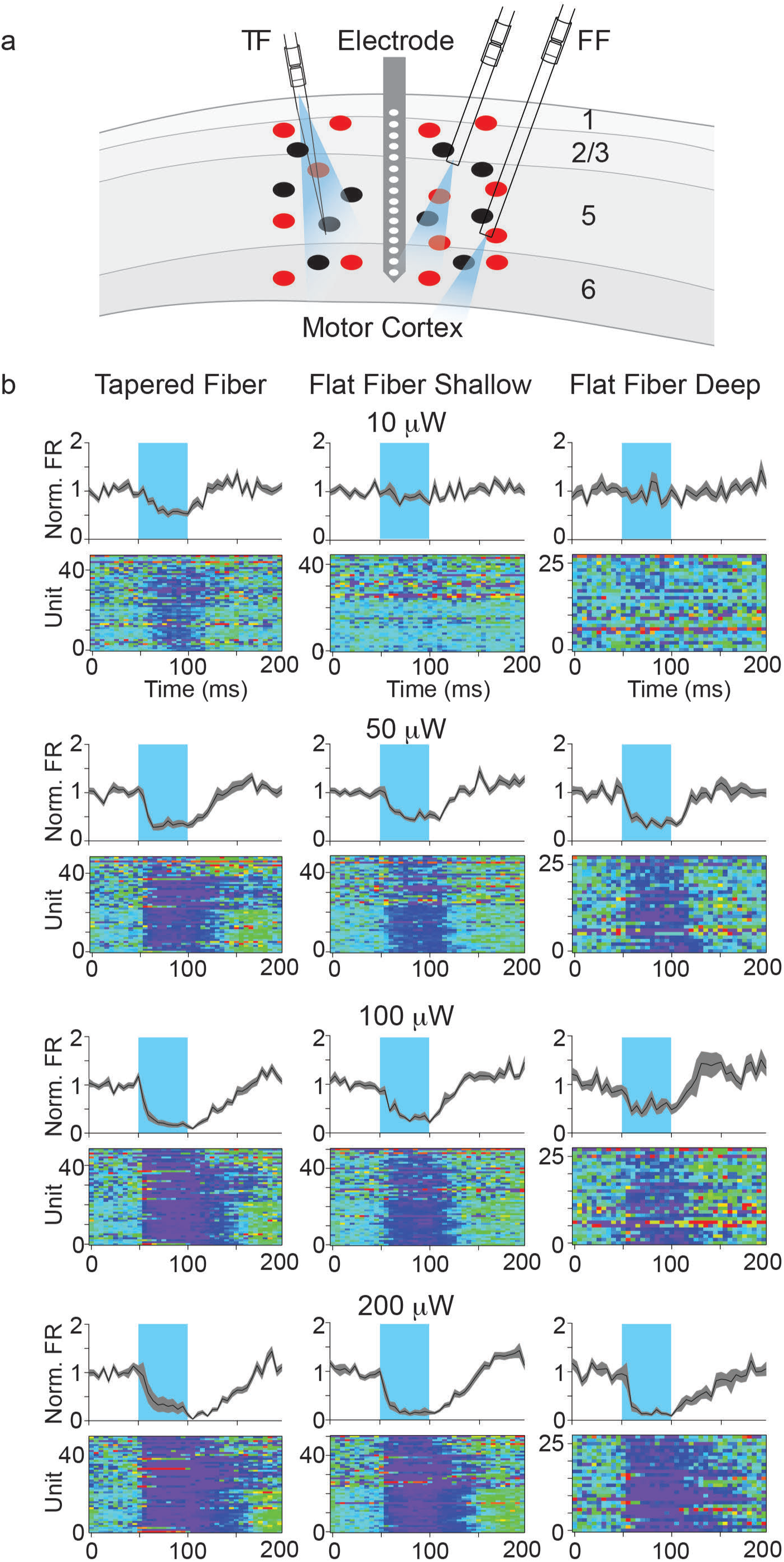
Optogenetic manipulation of motor cortex with TFs. **a**, Schematic of the experimental preparation. A 16 channel multi-electrode array is placed in primary motor cortex of a VGAT-ChR2 BAC transgenic mouse and maintained throughout the experiment. Units are recorded throughout cortical layers. A TF or FF in either a shallow or deep position is placed in cortex. The firing rates of individual units are compared during the basal period or during 50 ms optogenetic excitation cortical GABAergic interneurons **b**, Average normalized firing rates (black line with shaded area showing SEM) across cells with and without light (*t*op) and pseudocolored representations of normalized across-trial average firing rate of each unit as a function of time (*bottom*). The period of light delivery (50 ms) is shown in the cyan shaded regions. Data are shown for each fiber configuration (*left:* TF, *middle:* FF shallow, *right:* FF deep) and at 4 power levels (10, 50, 100, and 200 μW total emission from the fiber before implantation in the brain.

### Subsampling the volume of interest

A further benefit provided by TFs is the ability to dynamically control the illumination volume by changing the light input angle θ at the laser input end of the fiber. The angle θ defines the subset of guided modes injected into the fiber^13,14^ which in turn determines the cross section of the taper at which each is emitted. The position of the emitting section along the taper depends on θ as expected from the ray tracing simulations (Fig. S12 and Movie 1). In particular, low input angles allow light to outcouple close to the taper tip, whereas light injected at high input angles is mainly emitted at sections farther from the tip.

To characterize the geometrical emission properties of TFs as a function of θ (Fig. 5) we implemented a simple optical path in which θ is changed by translating a mirror (Fig. S13a). Monitoring the fluorescence generated by a TF inserted into a fluorescein solution shows that the emitting segment (~300 μm long) can be moved almost continuously along ~1 mm or ~1.5 mm, respectively, in 0.22 NA/ψ=2.2° or 0.39 NA/ψ=2.9° TFs (Fig. 5a,b). Importantly, total delivered light power is nearly independent on θ, apart for input angles very close to the maximum acceptance angle (Fig. 5c). To rapidly scan the illumination across brain volumes we used launching system with a scanning galvanometer and relay optics to change θ (Fig. S13b). This permits rapid switching between different emission segments (Movie 2) and near continuous movement of the emitting segment along the taper (Movie 3). Several launching paths can also be combined (Fig. S13c). This allows for multiple wavelengths to be outcoupled at the same time at different depths, with independently and addressable emission segments. This is shown in Movie 4 for simultaneous operation of a blue and a green light lasers.

**Figure 5.**
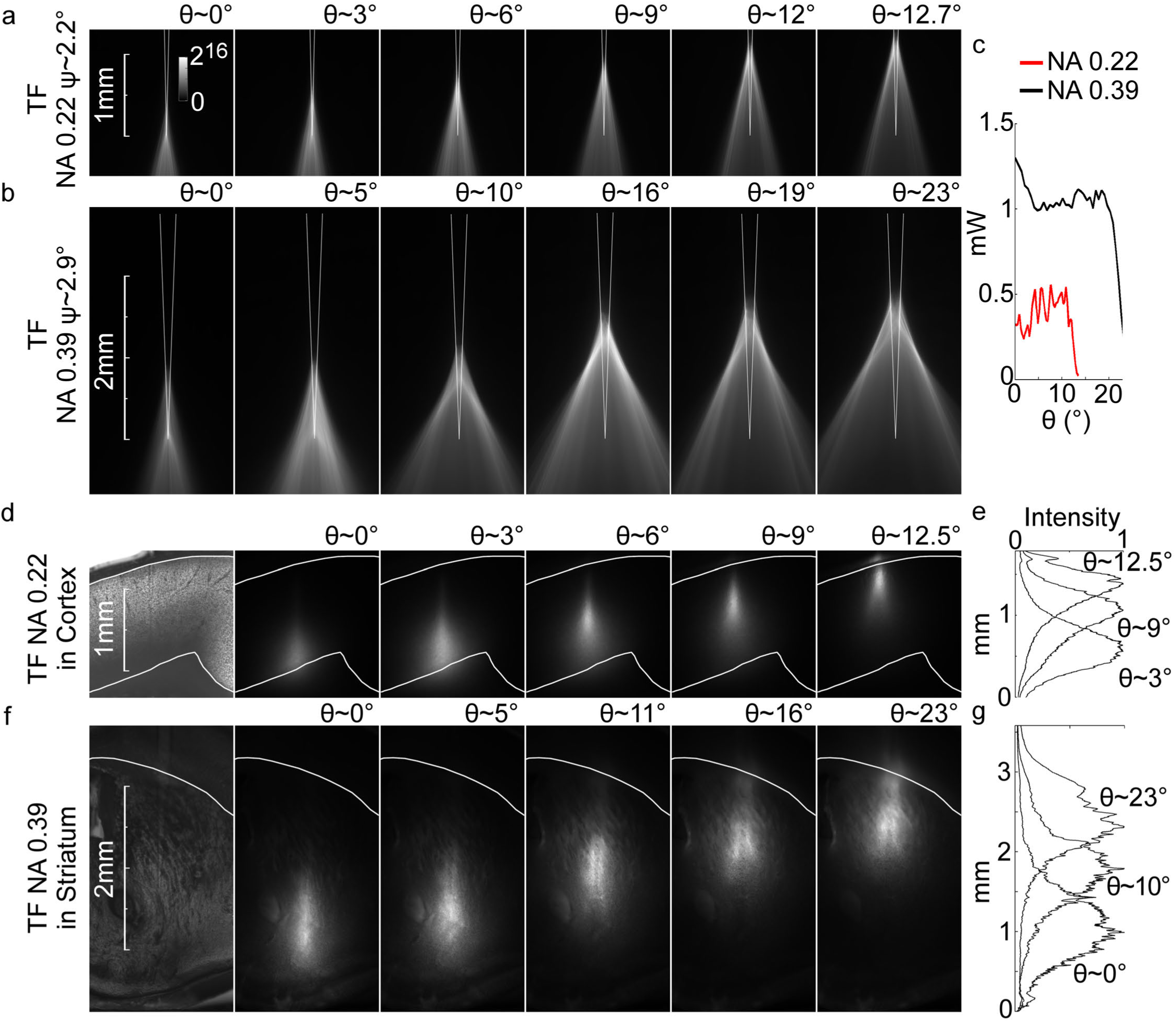
Site selective light delivery with TFs. **a**, Light delivery geometry for several values of light injection angle θ with a 0.22NA/ψ=2.2° TF into a fluorescein solution. Gray scale represents fluorescence intensity in arbitrary units and is the same for all panels. **b**, Light delivery geometry for several values of light injection angle θ with a 0.39 NA/ψ =2.9° TF into a fluorescein solution. **c**, Total Output power for a fixed input of 2.25 mW for TFs with 0.22 NA/ψ=2.2° TF (red line) and 0.39 NA/ψ=2.9° TF (black line) **d**, Site selective light delivery with a 0.22 NA/ψ=2.2° TF implanted into the cortical region of a fluorescently stained mouse brain slice. **e**, Normalized fluorescence intensity profiles, measured beside the taper, from the fluorescence images in (d). **f**, Site selective light delivery with a 0.39NA/ ψ=2.9° TF implanted into the striatum of a fluorescently stained mouse brain slice. **g**, Normalized fluorescence intensity profiles, measured beside the taper, from the fluorescence images in (f).

To examine the suitability of this technique for restricted light delivery in brain tissue, site-selective light delivery as a function of θ was evaluated in fluorescein-stained acute mouse brain slices. Both 0.22 NA/ψ= 2.2° and 0.39 NA/ψ = 2.9° TFs allowed near-continuous tuning of the illuminated brain region in both cortex and striatum (Fig. 5d–f). Tissue absorption and scattering shorten the propagation of emitted light into the tissue, further constraining the spatial geometry of the illuminated area. This leads to spatially separated light delivery volumes, resulting in an easy-to-use and versatile method to direct the light stimulus along a ~2 mm segment by implanting a single fixed fiber (Fig. 5e–g).

### *In vivo* multi-site stimulation

Selection of the emitting region of the taper is possible because different input angles corresponds to different sets of guided modes in the fiber^14^, which are outcoupled at different sections of the taper. However, while propagating into the fiber, a single subset of guided modes may undergo modal mixing induced by fiber impurities and bends. This could redistribute part of the guided light power to other modes, potentially resulting in a rearrangement of light emission along the taper. This possibility is particularly important for experiments requiring site-specific stimulation in moving animals that may induce unpredictable motions of the patch fiber. To evaluate the viability of using TFs for site-selective light delivery in moving mice, we measured the effects of fiber bending and shaking on light output from the TFs (Fig. S14). For a fixed input angle (θ=17°) the patch fiber carrying light to a TF with 0.39 NA/ψ = 2.9° was shaken and/or curved in order to simulate animal movement. Simultaneously, taper emission into a fluorescein droplet was recorded at high frame rate (~100 fps) with exposure time of 10 ms (Movies 5, 6 and 7). Light delivery fluctuations were quantified in terms of oscillations in the peak intensity, full-width at half maximum, and center of the light emission profile – fiber shaking and bending produced variations of all three metrics below 5% (Fig. S14).

To demonstrate the feasibility of multi-site optogenetic stimulation through a single TF in an individual animal, we examined spontaneous locomotion behavior of a mouse in an open arena (Fig. 6) using video recorded with depth time of flight cameras ^22^. TFs were designed (0.39 NA, ψ =2.3°) and implanted spanning the dorsal and ventral medial striatum of two mice (33.6g, and 31.7g at the time of implant) expressing ChR2 in all iSPNs as above (*Ador2a-Cre; Ai32*). An optical pathway was designed (Fig. 6a) and calibrated to deliver the same power of light from the distal (ventral striatum) and proximal (dorsal striatum) sites (Fig. 6c) using, respectively, 8° (θ_1_) and 22.5° (θ_2_) launch angles (Fig. 6a and Fig. S15).

**Figure 6.**
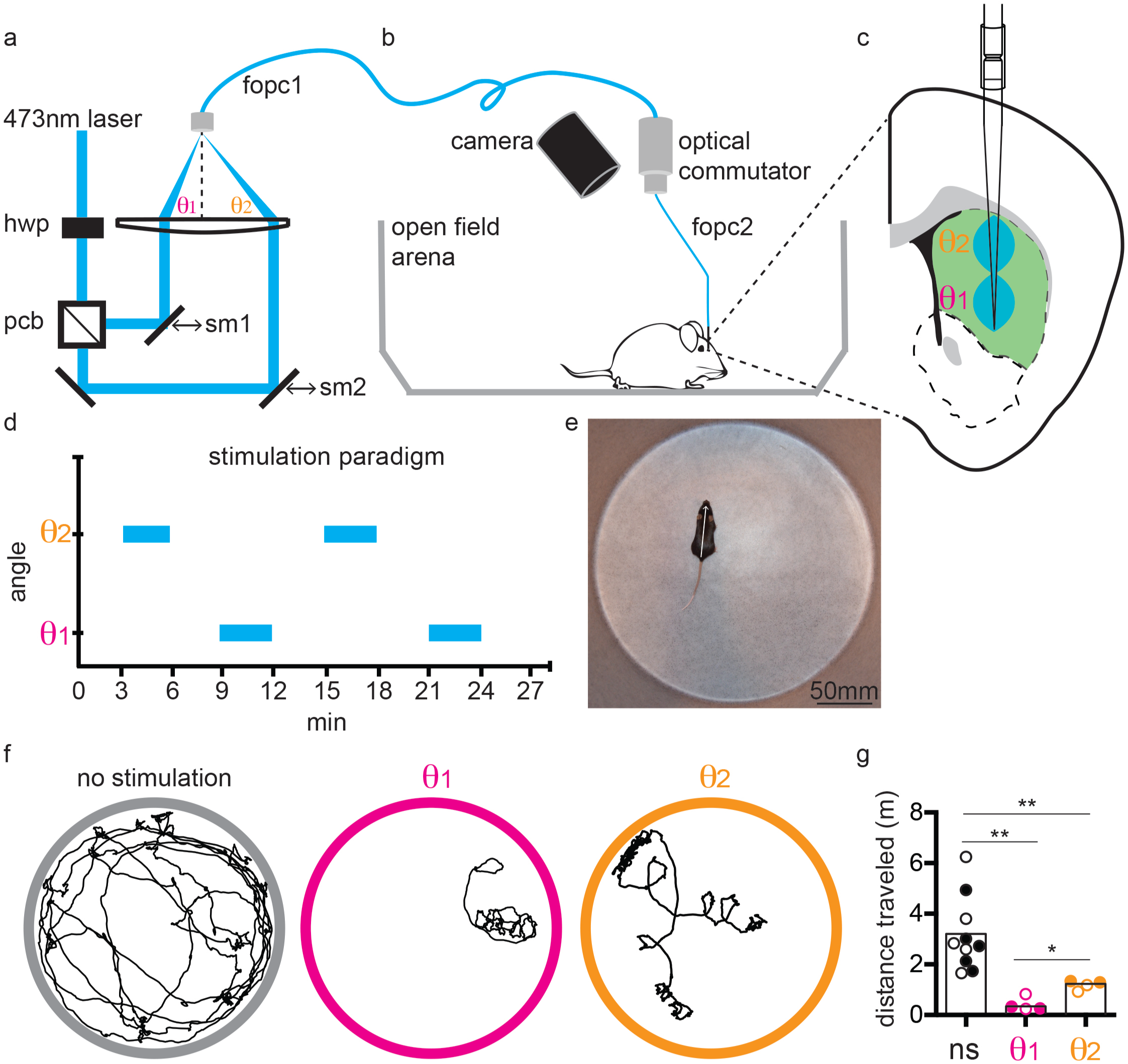
Selective light delivery with TFs in the open field. **a**, Schematic of optical setup. The output of a polarized laser was passed through a ½-wave plate (hwp) to rotate the polarization before entering a polarizing beam cube (pbc) that transmits and reflects, respectively, horizontally and vertically polarized light. Rotation of the hwp determines the fraction of laser light entering each path. Each of the laser paths is directed to a 2-inch collection lens via a sliding mirror (sm1 and sm2) that can be moved linearly to determine the launch angle into the first fiber-optic patch cord (fopc1). **b**, Fiber optic patch cable (fopc1) was connected from the optical pathway shown in (a) to a commercial optical commutator from which a second fiber optic patch cable (fopc2) led to the TF implanted in the animal. A camera above the arena monitored the location and depth of the mouse. **c**, The TF was implanted in the striatum (0.85A, 1.4L, 4.1D) of a transgenic animal expressing ChR2-YFP in iSPNs to allow for stimulation of two regions of striatum. Light input at θ1 (*magenta*) and θ2 (*orange*) angles resulted in emission that targeted the dorsal and ventral striatum, respectively. **d**, Example of stimulation paradigm in the open field arena on day 1. Animals spent a total of 27 min in the open field with 3 min sessions of either no light, light on at angle θ1, or light on angle at θ2. On a subsequent day of analysis the order of θ_1_ and θ_2_ stimulation were reversed. **e**, Snapshot of a mouse in the open field with an overlaid vector highlighting the simple feature extraction of position and orientation. **f**, Example of the positions of one mouse during 3-minute session of no stimulation (*left)*, dorsal stimulation (θ_1_, *middle*), and ventral stimulation (θ_2_, *right*). **g**, Quantification of distance traveled in the 3 conditions for the example mouse shown in (f). Significant differences were observed between the no stimulation (*left*), ventral stimulation (*middle*) and the dorsal stimulation (*right*) conditions. Bars indicate the means of all data points from individual 3 min blocks, which are shown by the circles (not filled: day 1; filled: day 2). *p < 0.05, **p<0.01, using a two-tailed Mann-Whitney U test.

Eight days after implant surgery, the mice were placed in an open circular arena and video recorded. Light was delivered to the brain via an optical commutator, a lightweight patch cord (200 μm Core, 0.39 NA, 1 meter long), and two fiber-fiber conjunctions as typically used for unrestrained mouse behavior experiments (Fig. 6a–b). The experimental paradigm consisted of 3 min blocks of either no stimulation (ns), or laser input to the fiber at angle 1 (θ_1_) or angle 2 (θ_2_) repeated in the following pattern ns θ_1_ ns θ_2_ ns θ_1_ ns θ_2_ ns, corresponding to alternating stimulation of ventral (θ_1_) and dorsal (θ_2_) striatum separated by periods of no stimulation (Fig. 6d). On the subsequent day of analysis the order of ventral (θ_1_) and dorsal (θ_2_) stimulation were reversed. Thus a total of 9 blocks per session were recorded in each of 2 days for the 2 mice. Basic video analyses of locomotion speed and orientation (Fig. 6e) reveal that stimulation at either dorsal or ventral striatum reduced locomotion and triggered contraversive spinning (Fig. 6f) with ventral stimulation via light injection at angle θ_2_ inducing more profound effects (distance traversed/3 min: ns: 3.16 ± 1.45m, n=10; ventral (θ_1_): 0.41m ± 0.27, n=4; dorsal(θ_2_): 1.18m ± 0.18, n=4; θ_1_ vs. θ_2_: p=0.0286. θ_1_ vs. ns: p = 0.002; θ_2_ vs ns: p =0.002) (Fig. 6g).

Mouse locomotion and posture (Fig. 7a) were analyzed using a machine learning approach that automatically detects repeated time varying “syllables” that correspond to the animal’s postural dynamics^22^. This technique produces a hidden Markov model in which each state encapsulates the postural dynamics of the mouse in each expressed syllable of behavior. The model is built using 3-dimensional postural information derived from each video frame collected (at 30 Hz) for both mice across all imaging sessions and stimulation conditions. In this case, 14 syllables were sufficient to explain on ~94% of locomotion behavior for the 2 mice in all sessions (ns: 87%; ventral (θ_1_): 98%; dorsal (θ_2_): 97%) (Fig. 7b). Furthermore, consistent with the previously described effects of iSPN stimulation on locomotion^23^ (Fig. 6) the most commonly observed behavior was pausing (syllable 1 and syllable 7) corresponding to a motionless mouse (in ns: 1.09%; ventral (θ_1_): 74%; dorsal (θ_2_): 38%) (Fig. 7b and Video 8) Within all the time included in these 14 syllables, the relative expression of the pause syllable varied across conditions, being most prominent in the ventral stimulation condition (Fig. 7c). Many of the remaining dominant non-pause, i.e. movement related syllables were also differentially expressed across stimulation condition (syllables 2-6 and 8-14: ns: 86%; ventral (θ_1_): 24%; dorsal (θ_2_): 59%) (Fig. 7d).

**Figure 7.**
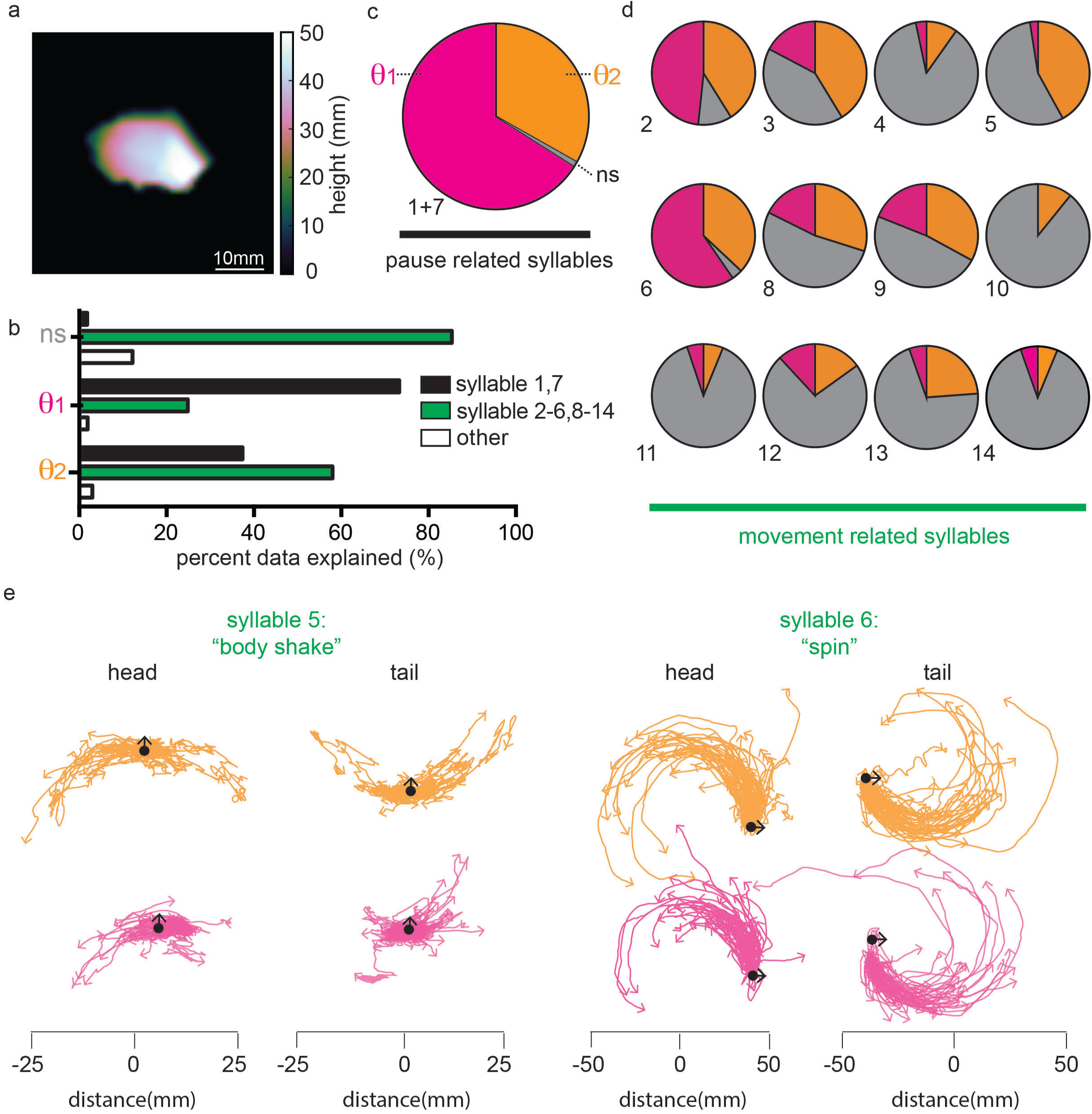
Mapping sub-second structure of behavior during optogenetic manipulation of ventral or dorsal striatum. **a**, Snapshot of a mouse in the open field with the height of the body at each pixel indicated by the color scale. **b**, The percent behavioral data explained in each condition (ns, θ_1_, and θ_2_ stimulation) by the major pause related (*black*) syllables (1 and *7*), the major movement-related (*green*) syllables (2-6 and 8-14) and all other syllables (*white*). **c**, Fractional expression of pause related syllables (1 and 7) during ns (*grey*) ventral (θ_1_) (*magenta*) and dorsal (θ_2_) (*orange*) conditions showing enhanced pausing with iSPN activation, particularly for ventral stimulation. **d**, Fractional expression of the dominant non-pause, i.e. movement related, syllables (2-6 and 8-14) color coded as in (c) showing differential expression of syllables across stimulation conditions. **e**, The trajectories of the head and tail of the mouse relative to its body center for syllable 5 (*left)* and syllable 6 (*right*) in either the ventral (θ_1_, *magenta*) and dorsal (θ_2_, *orange*) stimulation conditions. Data from 100 randomly chosen instances of each of syllables 5 and 6 expressed during ventral and dorsal stimulation are shown. The black dots depict the starting point of each of the aligned trajectories of the head and tail. The black arrows indicate the initial direction of head relative to the arena (i.e. not relative to the body center).

As described by these dominant 12 movement syllables, stimulation of ventral and dorsal striatum evoked different patterns of locomotion behavior. To demonstrate the difference in locomotion behavior between syllables as well as to highlight the consistency of behavior within a syllable across stimulation conditions, we plotted 100 instances of the head and tail trajectory relative to its body center in syllables 5 and 6 (Fig. 7e and Video 9 and 10).

## DISCUSSION

Here we demonstrate that TFs have substantial advantages for light delivery in the brain that might allow this tool to replace the flat-cleaved optical fibers that are typically used for optogenetic experiments. Three main features support this possibility. First, TFs are multipurpose such that the same device allows for either large-volume or site-selective light delivery. Second, by virtue of their tapered and smaller average cross section, TFs are minimally invasive and can be implanted directly into the brain region of interest. Third, they are simple to operate and compatible with optical equipment commonly present in neuroscience labs employing optogenetics. Although sharpened fibers have been sporadically used to increase light delivery angle or to reduce invasiveness^24,25^, their ability to provide uniform as well as dynamically controlled restricted illumination has not been previously recognized.

If used with standard fiber-coupled lasers or LEDs, TFs distribute light power more uniformly in the brain than do FFs. Indeed, light delivery is distributed over the taper surface and along nearly the entire linear extent of the fiber taper. This results in a better utilization of light power as it prevents the need to deliver too much light to cells close to the fiber face in order to stimulate more distant cells. Indeed, we find both more uniform stimulation of ChR2-expressing cells in a large brain volume (striatum) as well as more efficient silencing of a brain area cortex) by stimulation of inhibitory interneurons. Furthermore, the approach is viable for implantation in freely moving tethered mice as well as for acute insertion in head-fixed mice. The TF are also compatible with inexpensive LED-based commercial fiber launch systems (Fig. S16) which naturally inject light over a large NA and produce emission over the fiber face for large-volume stimulation.

When used to deliver light from the entire taper surface, TFs overcome one of the principal difficulties facing experiments in *in vivo* optogenetics – achieving uniform effective illumination of large brain structures with minimal invasiveness and light power. Although effective stimulation of neuron cell bodies expressing ChR2 can be obtained with power densities in the range 1-5 mW/mm^2^ [^26,27^] (and sometimes <1 mW/mm^2^, depending on the sensitivity of the ChR2-expressing cells)^28^, many published studies utilize orders of magnitude higher power^15,29–31^. This likely arises because in order to avoid excessive tissue damage, the FF is often positioned above the brain nucleus of interest. Due to the exponential fall off in power density from the FF, this necessitates the use of high light powers in order to stimulate cells throughout the nucleus^15,16,32^ and can result in local tissue heating in the range of several °C ^17^. By both permitting insertion of the fiber into the nucleus of interest and by delivering light along the linear extent of the taper, TFs can achieve efficient light stimulation. This feature may be even more important when considering inhibitory and spectrally shifted opsins that may require more light to achieve an effective neuronal perturbation. Nevertheless, for select applications, such as when the volume to be illuminated is extended in the orientation perpendicular that of the fiber, FF may provide a more effective light source.

Conversely, TFs also solve the problem of achieving on-the-fly adjustable delivery without moving the TF (Movies 1-3, Fig. 5–7), even of multiple light wavelengths (Movie 4) to sub-volumes of brain tissue. Site-selective light delivery is achieved using a simple optical setup that injects light into the fiber with a defined and adjustable angle θ. At the minimum, this setup consists of one lens and one translating mirror for slowly adjusting θ (Fig. S13a). For high-speed control, three lenses and one galvanometric mirror (Fig. S13b) are used to change θ quickly. With this approach, the emitting region can be modified step-by-step (every 20 ms in Movie 2) or continuously at various speeds (Movie 3).

From the perspective of an experimental neuroscientist, several classes of experiments become possible by exploiting the flexible and controllable nature of light delivery with TFs. For example, subregions of the striatum subserve different functions with specialized contributions to behavior being evident along the dorsal/ventral as well as medial/lateral axes and course topographic mapping to cortex laid out along the anterior/posterior axis^33^. This suborganization is difficult to access in a single experimental animal. As we demonstrate (Fig. 3,6–7), with a TF it is possible to illuminate an elongated column of striatum that spans the dorsal/ventral axis and compare, in a single animal, the effects of selective optogenetic manipulation of subregions along the fiber axis. Given that the striatum spans several millimeters in the mouse, these experiments are impractical with standard FF as these can neither deliver light to the entire structure nor to be repositioned for selective stimulation. Furthermore, with extended TFs a single fiber can be inserted to deliver light to both cortex and striatum. Since light of different wavelengths can be outcoupled from different zones and independently controlled (Movie 4), it will be possible to examine the effects of interactions between areas. For example, one can examine if the motor effects of inhibition of motor cortex can be overcome by excitation of the striatum.

In addition, as optogenetics becomes accessible in primates^34^, the larger brain structure will require the use of TFs for effective perturbations. For example, visual cortex of macaque is several millimeters in thickness with different cellular and receptive field properties in different layers ^35^. In this classic experimental system, current light delivery devices are unable to manipulate cells throughout all layers or, though a single device, test the effects of manipulations of superficial versus deep layer neurons. Furthermore, given the tight topographic organization of primate visual cortex and the use of individual animals for many recording sessions, it is valuable to have a device such as a TF with gentle taper for multiple insertions in each recording sessions.

In summary, we exploit TFs with small taper angles to achieve near uniform illumination of extended brain structures as well as to sub-sample regions of interest along a taper segment up to 2 mm. The devices were tested in both mouse motor cortex and striatum, showing ~5 times lower excitation power threshold and wider excitation volume, compared to standard fibers. Furthermore, we demonstrate their effective use through optical commutators and patch cords to compare the effects of stimulation of dorsal vs. ventral striatum in individual unrestrained mice spontaneously exploring an arena. Coupled to the minimum invasiveness of the device, the simplicity of the technique and its intrinsic compatibility with both laser and LED sources, we suggest that this approach can greatly complement existing methods for light delivery in optogenetics experiments and has the potential to replace commonly used flat-faced fibers for many applications.

## METHODS

### Ray tracing simulations

The commercial optical ray tracing software Zemax-OpticStudio (http://www.zemax.com/) was used to design and simulate the performances of TFs. The single TF was modeled as a straight core/cladding segment followed by a conical taper. The materials forming all the components of the TFs and the surrounding media were assumed homogenous (refractive index constant in space). The core/cladding diameters are 50/125μm and 200/225μm for fibers with numerical aperture NA=0.22 and NA=0.39, respectively. Core/cladding refractive indexes were set as specified by the fiber producer: 1.464/1.447 for NA=0.22 and 1.464/1.411 for NA=0.39. Since the tapers were obtained by heat and pull, commonly resulting in a melting of core and cladding materials in the tapered regions, taper refractive index was assumed as the average of core and cladding refractive indexes weighted to the core and cladding cross sectional areas. This resulted in tapers refractive index of 1.450 for fibers with NA=0.22 and 1.453 for fibers with NA=0.39. The length of the core/cladding block was set to 4mm, whereas the length of the taper is a function of the taper angle.

For results of simulations reported in Fig. 1b and Fig. S2, the source was modeled as a single ray injected into the fiber with a defined angle of incidence with respect to fiber optical axis. For simulations reported in Fig. 1 and Fig. S3 and S4 a monochromatic source (λ=473 nm) was modeled as a bundle of unpolarized parallel rays having a Gaussian intensity profile. The source was focused into the core/cladding section of the fiber through a Zemax model of the experimentally used aspheric condenser (Rochester Precision Optics www.rpoptics.com). Model A-375-A with focal length *f*=7.49 mm, NA=0.29, clear aperture of 4.50 mm was used for TFs with NA=0.22 (input Gaussian beam radius 1.66 mm measured at 1/e^2^, resulting in a focused waist of 1.6 μm RMS spot radius), whereas model A-390-A with *f*=4.60 mm, NA=0.47 and clear aperture of 4.90 mm was used for fibers with NA=0.39 (input Gaussian beam radius 1.84 mm measured at 1/e^2^, resulting in a focused spot of 1.9 μm RMS spot radius). The former coupler was used only with the NA=0.22 optical fiber, with a Gaussian beam radius of 1.66 mm which focuses on the fiber core with a 1.6 μm RMS spot radius; the latter was used with both fibers, with a Gaussian beam radius of 1.84 mm which produces a 1.9 μm RMS spot radius. For simulations displayed in Fig. S12 a parallel and unpolarized Gaussian beam with radius 25 μm at 1/e^2^ intensity was injected at different input angles.

The irradiance profile of the light out-coupled from the taper was recorded through a rectangular pixelated detector laid along the taper sidewall (Figure S4). Detector length was set as the taper side length and its width to 20 μm. These profiles are then averaged along the short side of the detector and fitted with a Gaussian function. The full-width at half maximum criterion was used to retrieve the emission length (*L_0.5_*, measured axially from the taper tip) from the irradiance profiles. For each single ray tracing session, 5M rays were launched into the system and each ray was split at the boundary between two different media, according to Fresnel coefficients.

### TFs stub preparation and optical measurements

Tapered optical fibers with taper angles in the rage 2°< ψ <8° were obtained from OptogeniX (www.optogenix.com) and connected to a ceramic ferrules following the procedure described previously^36^. A ceramic ferrule-SMA patch cable was used to connect the fiber stub to the optical setup displayed in Fig. S4 for full NA light injection or to the setups schematized in Fig. S13 for site-selective light delivery. The taper was immersed into a fluorescent water:fluorescein solution or inserted in stained brain slices positioned under a 5X objective of a fluorescence Zeiss microscope equipped with a FITC filter. Images were acquired with a Hamamatsu Orca Flash 4.0. sCMOS camera at a resolution 2048x2048 pixel (pixel depth 16 bit). Optical output power was measured in air. In the case of the multiwavelength experiment reported in Movie 4, the taper was immersed into a drop of diluted milk to induce scattering of outcoupled light and detected with a color CCD camera.

For imaging light delivery geometry in fixed brain slices, slices of thickness ranging from 200 μm to 400 μm were permeabilized with Triton X-100 0.1% for 10 min to allow homogenous cell staining through the whole brain slice thickness and then washed with 1X PBS three times. Slices were then incubated with Sybr Green 1:10.000 (Thermo Fischer Scientific Inc.) for 20 min on an orbital shaker in dark and thoroughly washed with 1X PBS.

### Multielectrode array recordings

All mouse handling and manipulations were performed in accordance with protocols approved by the Harvard Standing Committee on Animal Care following guidelines described in the US National Institutes of Health Guide for the Care and Use of Laboratory Animals. Recordings in primary motor cortex and off-line analysis of spiking rates were accomplished as previously described ^18^. Mice were habituated to head-restraint prior to the recording session. Only one fiber optic (TF or FF) was inserted at a time and the effects of transient stimulation of cortical inhibitory neurons on identified units were examined. For each unit, spiking rates were normalized to the pre-illumination rate. Averages were calculated across neurons. Two mice were used for these experiments and both were B6.Cg-Tg(Slc32a1-COP4*H134R/EYFP)8Gfng/J purchased from Jackson Laboratory (Stock #014548).

### Analysis of c-fos induction

Transgenic adult male animals (weight 26-42 grams) were anesthetized with isoflurane and placed in a small animal stereotaxic frame (David Kopf instruments). Under aseptic conditions, the skull was exposed and a small hole was drilled. Animals received 0.01mg/gram of sterile Ketofen (Zoetis). For FF implanted animals and the lateral striatum TF implanted animal the hole was made at 0.85mm anterior, 1.95mm lateral from bregma. For the medial striatum TF implanted animals the hole was made at 0.85mm anterior, 1.65mm lateral from bregma. A TF (3.7mm from pia) or FF (2.3 mm from pia) was inserted manually using a cannula holder (David Kopf instruments). The TF or FF was then glued in place (454 instant adhesive, Loctite). And the skull was covered with dental cement (CandB Metabond, Parkell inc). Regardless of the implant-surgery time animals were under isoflurane anesthesia for a total of 1 hour before stimulation started. Stimulation was delivered in 30-second on/off cycles for a total of 1 hour (473 nm, 1 mW outputted at fiber exit). 2 hours post stimulation animals were euthanized with 0.2 ml of 10% Fatal Plus solution (Vortech pharmaceuticals) in saline before being perfused with 4% paraformaldehyde (PFA) in 0.1M sodium phosphate buffer (PBS). Brains were post fixed for 24 hours in PFA, washed in PBS, and sectioned (50um) coronally.

Immunohistochemistry conditions were the same for all animals. In short, slices were incubated in PBS blocking solution containing 0.3% TritonX (PBST) for 1 hour at RT. Slices were then incubated over night at 4°C in the same blocking solution with 1% goat serum and 1ug/ml c-Fos rabbit polyclonal IgG antibody (H-125, Santa Cruz Biotechnology). The next morning slices were rinsed 3x10min in PBS before being incubated in the blocking solution with secondary antibody (1mg/ml goat anti-rabbit Alexa Fluor 647 or Alexa Fluor 594, Life Technologies). The slices were then rinsed again and mounted. After drying, slices were coverslipped with ProLong antifade mounting media containing DAPI (Molecular Probes) and imaged with an Olympus VS 120 slide-scanning microscope using a 10x objective.

### Open field surgery

The surgery for TF implant was as described above with coordinates for the TF in striatum being 0.85A, 1.4L, 4.1D. Animals recovered for 5 days post surgery and were handled for 3 days before experimentation.

### Behavior

On the day of the experiment animals were positioned in the open field arena. The experimental paradigm consisted of 3 min blocks of either no stimulation (ns), or laser input to the fiber at angle 1 (θ_1_) or angle 2 (θ_2_) repeated in the following pattern ns θ_1_ ns θ_2_ ns θ_1_ ns θ_2_ ns, corresponding to alternating stimulation of ventral (θ_1_) and dorsal (θ_2_) striatum separated by periods of no stimulation. On the subsequent day of analysis the order of ventral (θ_1_) and dorsal (θ_2_) stimulation were reversed.

### Open field analysis

Mice were recorded using a Microsoft Kinect (v1), which records depth video data at 30 frames per second. For data in figure 6, scalar features were extracted using previously published methods^22^.

For modeling data in figure 7, the open field behavior was analyzed using previously published methods^22^. In brief, the data were subjected to machine learning methods that describe the mouse’s behavior as re-usable sub-second modules, or syllables. All free parameters were set to the values described in Wiltschko et al^22^ with the exception of the stickiness parameters, kappa, which sets the model’s tendency to remain in the same syllable over time (rather than switch between different syllables). This parameter was tuned so that the overall syllable duration distribution qualitatively matched a model-free analysis of behavioral change-points as in Wiltschko et al^22^. For this analysis we set kappa to 291600. For head and tail positioning of the mouse in Fig. 7, the head and tail position were computed first by fitting an ellipse to the image of the mouse^22^. Then the head and tail positions were defined as the two farthest points of the ellipse (i.e. the points of the ellipse that intersect with its principal axis).

### Data availability

All data and code are available from the authors.

